# Genomic Perspectives on the Emerging SARS-CoV-2 Omicron Variant

**DOI:** 10.1101/2022.01.05.474231

**Authors:** Wentai Ma, Jing Yang, Haoyi Fu, Chao Su, Caixia Yu, Qihui Wang, Ana Tereza Ribeiro de Vasconcelos, Georgii A. Bazykin, Yiming Bao, Mingkun Li

## Abstract

A new variant of concern for SARS-CoV-2, Omicron (B.1.1.529), was designated by the World Health Organization on November 26, 2021. This study analyzed the viral genome sequencing data of 108 samples collected from patients infected with Omicron. First, we found that the enrichment efficiency of viral nucleic acids was reduced due to mutations in the region where the primers anneal to. Second, the Omicron variant possesses an excessive number of mutations compared to other variants circulating at the same time (62 *vs*. 45), especially in the *Spike* gene. Mutations in the *Spike* gene confer alterations in 32 amino acid residues, which was more than those observed in other SARS-CoV-2 variants. Moreover, a large number of nonsynonymous mutations occur in the codons for the amino acid residues located on the surface of the Spike protein, which could potentially affect the replication, infectivity, and antigenicity of SARS-CoV-2. Third, there are 53 mutations between the Omicron variant and its closest sequences available in public databases. Many of those mutations were rarely observed in the public database and had a low mutation rate. In addition, the linkage disequilibrium between these mutations were low, with a limited number of mutations (6) concurrently observed in the same genome, suggesting that the Omicron variant would be in a different evolutionary branch from the currently prevalent variants. To improve our ability to detect and track the source of new variants rapidly, it is imperative to further strengthen genomic surveillance and data sharing globally in a timely manner.

## Introduction

On November 22, 2021, the first genome sequence of a new variant of concern (VOC), Omicron (also known as B.1.1.529), was released in GISAID (Global initiative on sharing all influenza data) (EPI_ISL_6590782) [1]. The sample was obtained from a patient who arrived in Hong Kong on November 11 from South Africa via Doha in Qatar (https://news.sky.com/story/covid-19-how-the-spread-of-omicron-went-from-patient-zero-to-all-around-the-globe-12482183). To date, the first known Omicron variant sample was collected on November 5, 2021 in South Africa (EPI_ISL_7456440). Until December 12, 2021, there were over 2000 Omicron sequences submitted to the GISAID from South Africa, Botswana, Ghana, the United Kingdom, and many other countries. The emergence of this variant has attracted much attention due to the sheer number of mutations in the *Spike* gene, which may affect the viral transmissibility, replication, and binding of antibodies, and its dramatic increase in South Africa [2]. Preliminary studies showed that the new variant could substantially evade immunity from prior infection and vaccination [3,4]. Meanwhile, a preprint report proposed that the emergence of the Omicron variant was associated with an increased risk of SARS-CoV-2 reinfection [5]. However, it is still unclear where the new variant came.

In this study, we characterized the genomic features of the Omicron variant using data from 108 patients infected with the Omicron variant, which were generated by the Network for Genomic Surveillance in South Africa (NGS-SA) [2,6], and we speculate that the new variant is unlikely derived from recently discovered variants through either mutation or recombination.

## Results

### Reduced enrichment efficiency of the PCR-tiling amplicon protocols on the Omicron variant

Of the 207 Omicron samples sequenced, 158 samples had more than 90% of the viral genome covered by at least 5-fold, which were used in the subsequent analysis. Notably, two sequencing protocols were implemented. The first was to enrich the viral genome with the Midnight V6 primer sets followed by sequencing on the GridION platform (hereinafter referred to as Midnight, dx.doi.org/10.17504/protocols.io.bwyppfvn). The second protocol involved enrichment by the Artic V4 primer set, and the amplicons were sequenced on the Illumina MiSeq platform (hereinafter referred to as Artic, dx.doi.org/10.17504/protocols.io.bdp7i5rn). Fifty samples were sequenced using both protocols, and we found a high consistency in the major allele frequency between the two protocols (Figure S1). Artic data were preferred due to higher sequencing depth (median: 191 *vs*. 250, *P* < 0.01, Mann–Whitney U test). Finally, 49 samples sequenced by the Midnight protocol and 59 samples sequenced by the Artic protocol were included in the study.

Both protocols enabled efficient enrichment of viral nucleic acids from total RNA, the fraction of SARS-CoV-2 reads in the sequencing data were 84% and 94%, respectively for the Midnight and the Artic protocol. Although the Artic protocol had a relatively higher in-target percentage (*P* < 0.001, Mann–Whitney U test), the evenness of the sequencing depth of the SARS-CoV-2 was higher for the Midnight protocol (variance of the sequencing depth, 0.121 *vs*. 0.159, *P* < 0.001, Mann–Whitney U test). The sequencing depth profile of the SARS-CoV-2 genome was similar among samples sequenced by the same protocol but differed markedly between the two protocols (**Figure 1**A). The sequencing depth varied among different genomic regions, reflecting the differential enrichment efficiency of the primers. Moreover, we found that the large number of mutations possessed by the Omicron variant had a significant impact on the efficiency of the primers. In particular, seven primers in the Artic protocol were affected by at least one mutation, and three primers in the Midnight protocol were affected (Figure 1A). The worst coverage of the three regions for Primers 76, 79, and 90 using the Artic protocol were all associated with the presence of mutations in the region where these primers annealed to, whose sequencing depths were reduced by 2586, 246, and 234-fold, respectively, compared to the expected depth (Figure 1B). Strikingly, five mutations were located at the 5’ end of the least efficient Primer 76. The enrichment efficiency of other four primers (Primers 10, 27, 88, 89) was less affected by the mutations, which showed 1.3, 1.4, 3.4, and 1.9-fold reductions, respectively. Thus, the results suggest that the Omicron mutations can decrease the enrichment efficiency by PCR amplification, and there is an urgent need to update the Arctic V4 primers. We noted that the developer of the Artic protocol had already proposed a solution on this, and all seven affected primers had been updated (https://community.artic.network/t/sars-cov-2-v4-1-update-for-omicron-variant/342). In contrast, the efficiency of Midnight primers was less influenced by mutations in the Omicron variant. The three affected primers, Primers 10, 24, 28, showed no reduction, 2-fold, and 28-fold reduction respectively in sequencing depth compared to the expected depth.

**Figure 1.**
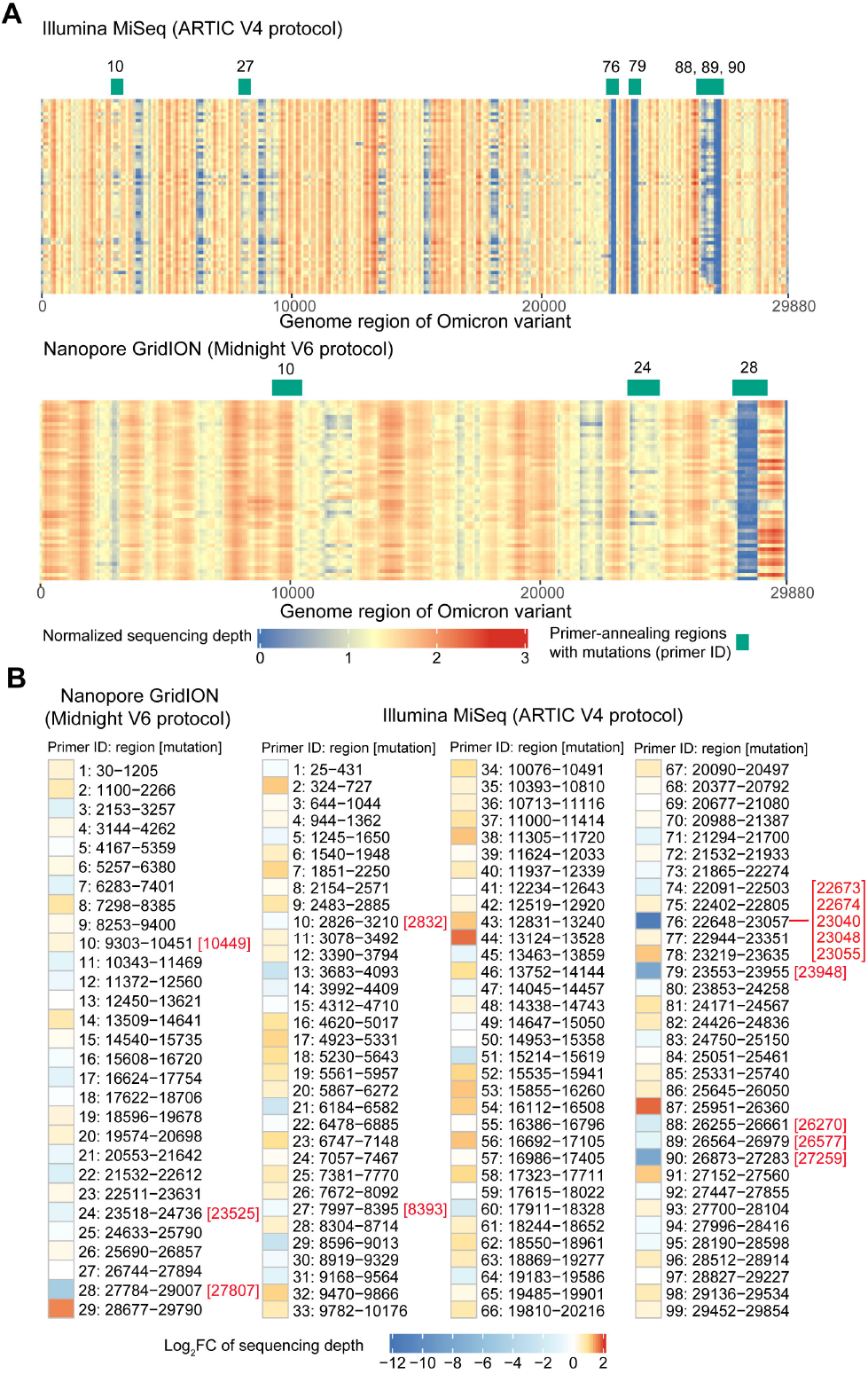
Sequence enrichment efficiency of the Omicron variant using different protocols. **A**. Distribution of the sequencing depth of the Omicron variant. The average sequencing depth is shown for each non-overlapping window of 100 bp after normalization by the total number of reads in the sample. The primers affected by the mutations in Omicron are labeled on top of the figure. **B**. The efficiency of each primer in amplifying the nucleic acids of the Omicron variant. The color represents the fold change of enrichment efficiency, calculated by the sum of the depth of all samples in this region divided by the expected value (assuming no differences among regions). The overlapping region of adjacent primers was excluded from the analysis. The Omicron mutations that located in the region where primers anneal to are labeled on the right of the primer ID.

### An extraordinary number of mutations in the *Spike* gene of the Omicron variant

The number of mutations (with major allele frequency ≥ 70%) of the Omicron variant varied from 61 to 64, and 61 of them were identified in more than 90% of the samples, which included 54 SNPs, six deletions, and one insertion. All these mutations were fixed at the individual level (**Figure 2**A). The total number of mutations was significantly higher than that of other variants detected in South Africa in November (median 62 vs. 45, *P* < 0.001, Mann–Whitney U test). Strikingly, over half of these mutations (34, 55.7%) were located in the *Spike* gene, whose length was 12.8% of the whole genome. Moreover, 32 of these mutations were nonsynonymous mutations. The proportion was significantly higher than that observed in the same region in other variants (94% vs. 67%, *P* < 0.001, Fisher’s exact test, Ka/Ks [7] = 8.65), suggesting positive selection on this gene.

**Figure 2.**
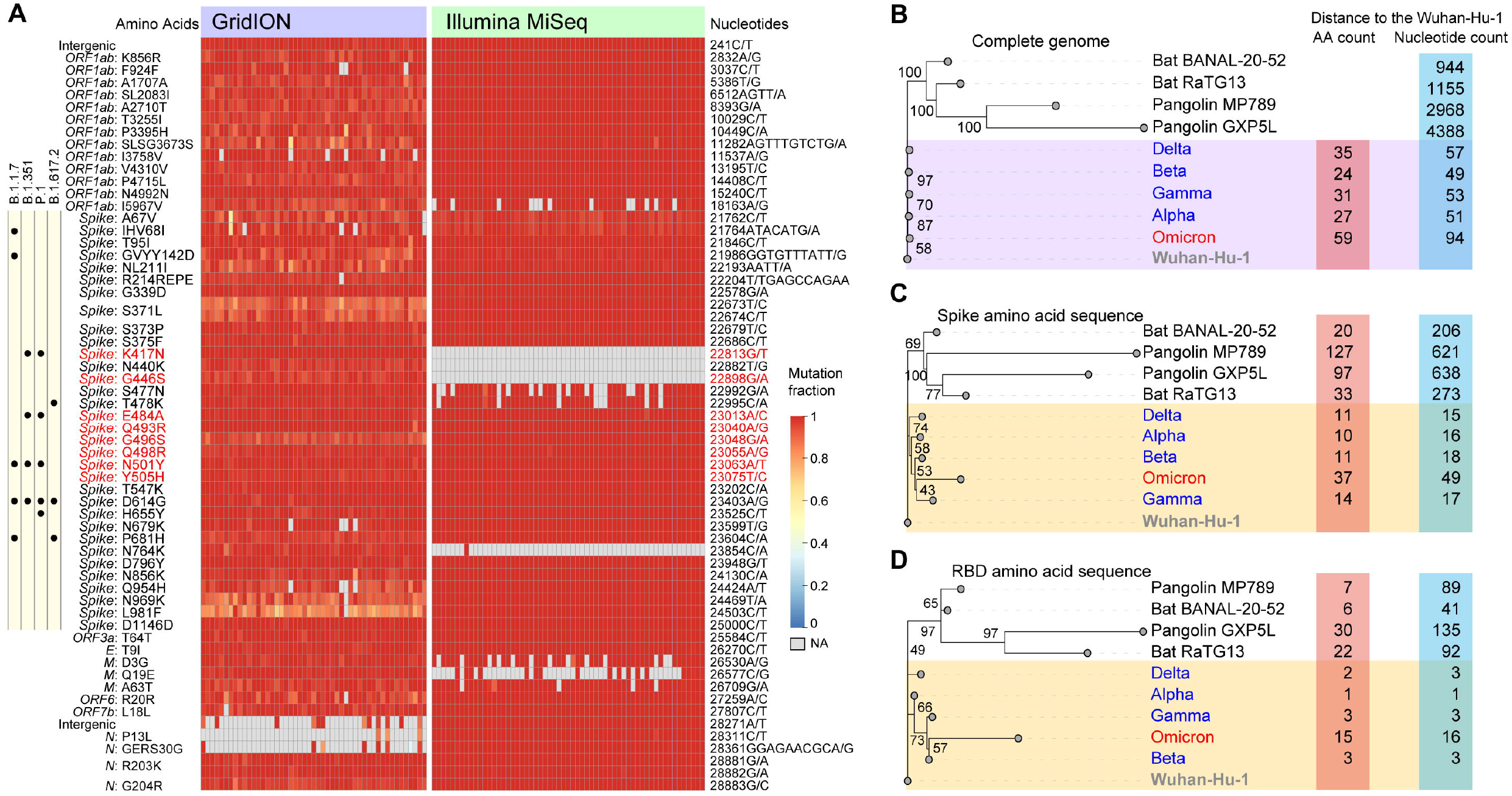
Mutations in the Omicron genome and its evolutionary relationship with other variants and SARS-like coronaviruses. **A**. Summary of mutations in the Omicron genome. Each row represents a mutation, and changes in nucleotides and amino acids are marked on two sides of the heatmap. Mutations located in the sites critical for viral binding to the human receptor angiotensin-converting enzyme 2 (ACE2) are marked in red [13]. Mutations observed in the *Spike* gene of other variants of concern (VOCs) are listed on the left of the heatmap. **B**. Phylogenetic tree of five VOCs and SARS-like coronaviruses based on the nucleotide sequences. **C**. Phylogenetic tree of five VOCs and SARS-like coronaviruses based on the amino acid sequences in the *Spike* gene. **D**. Phylogenetic tree of five VOCs and SARS-like coronaviruses based on the amino acid sequences in the RBD region. Two bat coronaviruses (Bat BANAL-20-52 and Bat RaTG3) whose genomes were most similar to SARS-CoV-2 [8,9], two pangolin coronaviruses (Pangolin MP789 and Pangolin GXP5L) [10,11], and sequences of other recently collected VOCs (EPI_ISL_6141707, EPI_ISL_6774033, EPI_ISL_6898988, EPI_ISL_6585201 for the Alpha, Beta, Gamma, and Delta variants, respectively. All sequences were collected in November 2021, and those collected in South Africa were preferred) were included in the analysis of the phylogenetic tree. The Wuhan-Hu-1 sequence is shown as the outgroup of the tree for better visualization [12]. The number of mutations relative to Wuhan-Hu-1 is listed on the right of the tree. Insertions or deletions of multiple bases were considered as a single mutation.

The Omicron variant showed a greater number of mutations than other VOCs (Figure 2B). The difference was more marked in the *Spike* gene, where the Omicron variant possessed 2-15 times more amino acid changes than other VOCs collected simultaneously (Figures 2C, D). Strikingly, the divergence in the amino acid sequence between the Omicron variant and the early SARS-CoV-2 sequence (Wuhan-Hu-1) in the *Spike* and RBD regions was greater or equivalent to that between SARS-like coronavirus (Pangolin MP789, Bat BANAL-20-52, and Bat RaTG13) and Wuhan-Hu-1 [8–12]. The dramatic changes in the *Spike* and RBD regions may substantially change the antigenicity and susceptibility to pre-existing antibodies.

### Potential risks associated with Omicron mutations

Most mutations occurred on the surface of the trimeric spike protein, especially in the RBD region (Figure S2). Eight of the 16 mutations in the RBD (K417N, G446S, E484A, Q493R, G496S, Q498R, N501Y, and Y505H) were located at positions that were proposed to be critical for viral binding to the host receptor angiotensin-converting enzyme 2 (ACE2) [13]. Among them, the K417N and N501Y mutations, which were also identified in the Beta variant, were reported to influence binding to human ACE2 [14]; N501Y confers a higher affinity of the viral Spike protein to ACE2 [15]. How the other mutations affect the affinity to ACE2 of humans and other animal hosts is still unknown.

Moreover, some other mutations in the *Spike* gene are known to be associated with changes in replication and infectivity of the virus. For example, Δ69-70 could enhance infectivity associated with increased cleaved Spike incorporation [16]; P681H could potentially confer replication advantage through increased cleavage efficacy by furin and adaptation to resist innate immunity [3,17]; H655Y was suspected to be an adaptive mutation that could increase the infectivity of the virus in both human and animal models [16]. In addition, mutations in other genes, such as R203K and G204R in the *Nucleoprotein* gene, could also potentially increase the infectivity, fitness, and virulence of the virus [18]. Of note, the function of these mutations was investigated because they were present in other VOCs. The effect of other less frequent mutations and the combination of the aforementioned mutations on the biology of the virus warrants further investigation.

Mutations in the RBD region, which is the target of many antibodies, may compromise the neutralization of existing antibodies induced by vaccination or natural infection [19]. Recent studies have shown severely reduced neutralization of the Omicron variant by monoclonal antibodies and vaccine sera [4,20,21]. Meanwhile, preliminary studies suggested that the Omicron variant caused three times more reinfection than previous strains, further supporting the speculation that the new variant can evade immunity from prior infection and vaccination [5]. However, the escape was incomplete, and a vaccine booster shot is likely to provide a high level of protection against the Omicron variant [4]. Here, we analyzed the epitope regions of 182 protein complex structures of antibodies that bound to SARS-CoV-2 Spike, the RBD, or NTD from the Protein Data Bank. We found that mutations in the Omicron variant were enriched in the epitope region of the Spike protein (**Figure 3**A). The median number of antibodies bound to the Omicron mutation sites was 53, which was significantly higher than those bound to other positions (median 3, *P* < 0.001, Mann–Whitney U test). Moreover, we found that these mutations could potentially impact the binding of different classes of antibodies (by analyzing the deep mutational scanning data [22], Figure 3B), which was classified by the location and conformation of antibody binding [23], suggesting that the therapeutic strategy of antibody cocktails may also be affected.

**Figure 3.**
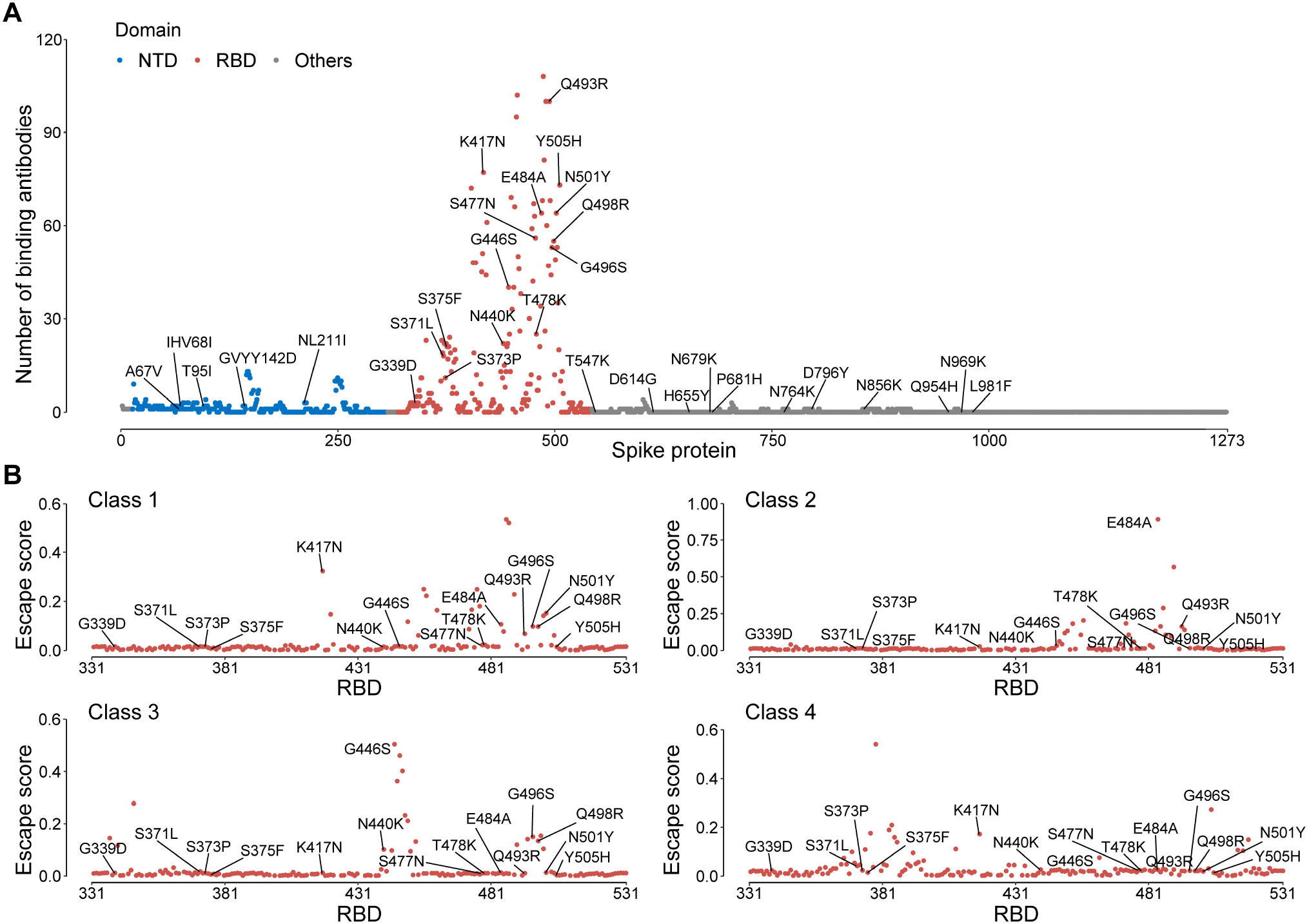
Distribution of the Omicron mutations at the antibody binding positions. **A**. The number of binding antibodies at the Omicron mutation sites. **B**. The escape score of the Omicron mutations estimated from deep mutational scanning. The escape score for each position was calculated as the mean of the scores of all antibodies belonging to the same class. All Omicron mutations were labeled on the figure.

### Obscure evolutionary trajectory of the Omicron variant

In addition to the 61 shared mutations, some private mutations were identified in different individuals, ranging from one to three, indicating relatively low population diversity at the time of sampling (**Figure 4**A). Meanwhile, no obvious clusters were found in the phylogenetic tree, suggesting that the Omicron variant was still in the early transmission stage during sampling. The time to the most recent common ancestor (TMRCA) was estimated to be in the middle of October 2021 (95% highest density interval: October 7 to 20).

**Figure 4.**
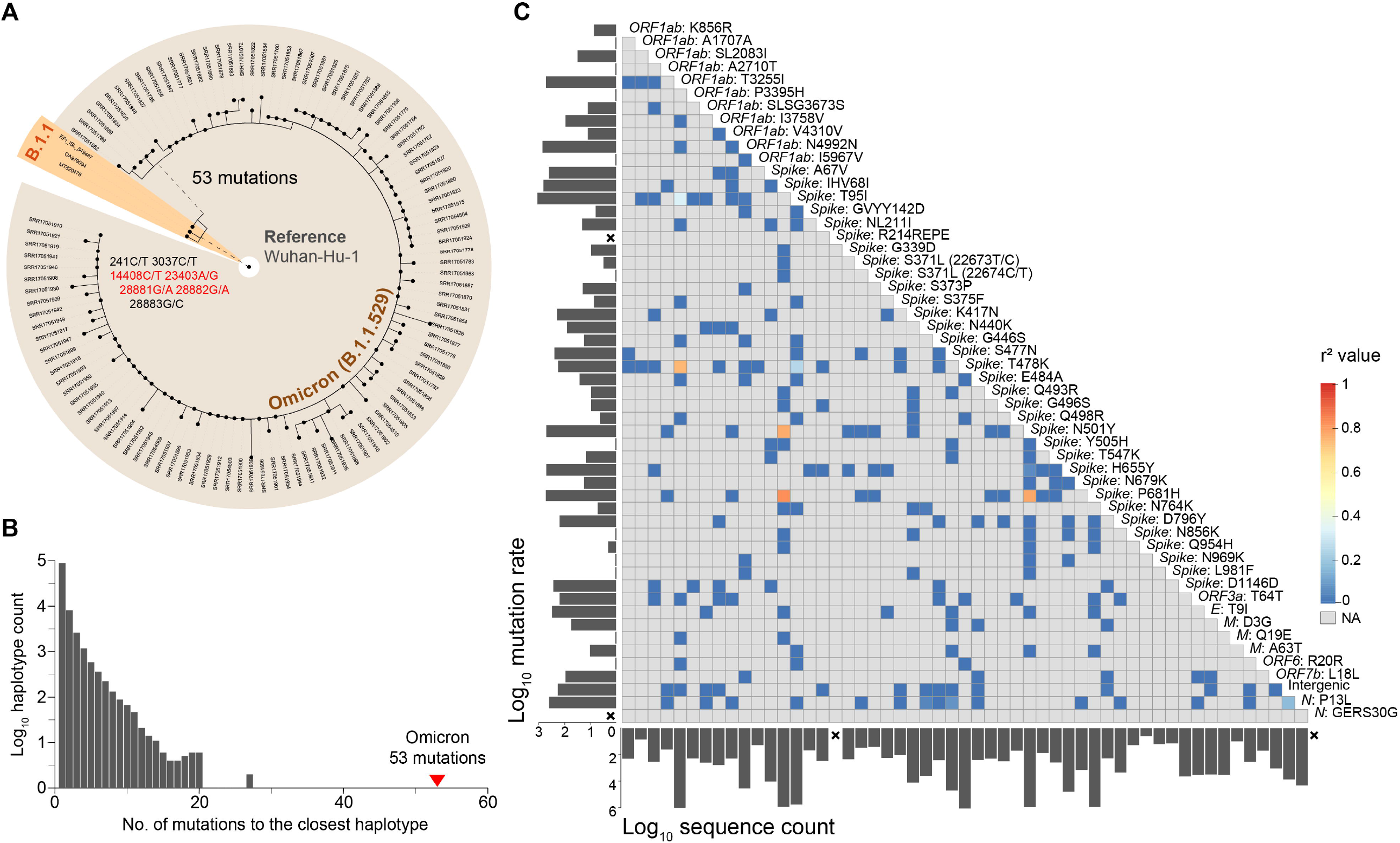
Evolutionary features of the Omicron variant. **A**.The phylogenetic tree of 108 Omicron sequences and their closest sequences in the database. Wuhan-Hu-1 is shown as the outgroup of the tree. The three closest sequences belonging to lineage B.1.1 are highlighted in orange. Nonsynonymous mutations are marked in red. **B**. The distribution of the number of differences between all haplotypes (nonredundant sequences) in the public database and their closest sequences. The minimum number of sequences required for a valid haplotype was set to 3. **C**. Correlation between different Omicron mutations. Only 54 Omicron lineage-specific mutations were included in the analysis. The color in the heatmap represents the linkage disequilibrium coefficient (*r*^*2*^) between mutations. The mutation rate and the number of sequences in the public database that possess the same mutation are labeled on the left and bottom of the heatmap, respectively. A cross is labeled if the mutation was not observed in the public database.

To screen for the possible predecessors of the Omicron variant, the 108 Omicron sequences were used as queries to look for the closest sequences in the public database, which included more than 5 million sequences released before November 1, 2021. We found three closest sequences to the queries, which differed by 53-56 nucleotides from the Omicron genomes. The three sequences were from lineage B.1.1 and collected between March and June 2020. They all had eight mutations relative to Wuhan-Hu-1, and seven of the mutations were shared among them (Figure 4A). The large number of differences suggests that the Omicron lineage was separated from other lineages a long time ago and has never been sequenced since then. This is an uncommon situation considering more than 5 million genomes have been sequenced in over 180 countries and regions. The distribution of the number of differences between all sequences in the public database and their closest sequences showed that 53-56 is approximately three times higher than the maximum number of differences observed in the database (20 when at least three sequences were required to eliminate the influence of sequencing or assembly errors, Figure 4B), again emphasizing the distinctiveness of the Omicron variant.

Most Omicron lineage-specific mutations (52/54) were identified in public databases (Figure 4C). However, they were unlikely to be presented in one sequence by chance. First, over half of the mutations were rarely detected in the populations, i.e., 33 mutations were detected in less than 1000 samples out of five million sequences (16 mutations were detected in less than 100 samples). Second, the mutation rate (represented by the occurrence number of mutations on the phylogenetic tree) was extremely low for 13 of the mutations (occurring only once in the evolution of SARS-CoV-2, mutation rate = 1). Third, the linkage disequilibrium between these mutations was low, and only four mutation pairs had *r*^*2*^ greater than 0.8. Moreover, we further examined whether any combination of these mutations appeared in the database and found that the maximum number of mutations in the same genome was six. Therefore, the evolutionary trajectory of the Omicron lineage cannot be resolved by the current genome data.

## Discussion

The unique genome features of the Omicron variant make it the most special SARS-CoV-2 variant to date. The excess number of nonsynonymous mutations in the Spike gene implies that the Omicron variant might evolve under selection pressure, which may come from antibodies or adaptation to new hosts. It is speculated that it may have been incubated in a patient chronically infected with SARS-CoV-2, e.g., HIV patients with immunocompromising conditions. This hypothesis was supported by the accelerated viral evolution observed in immunocompromised patients and has been previously proposed to explain how the Alpha variant was generated [24,25] (https://virological.org/t/preliminary-genomic-characterisation-of-an-emergent-sars-cov-2-lineage-in-the-uk-defined-by-a-novel-set-of-spike-mutations/563). If this hypothesis is true for the Omicron variant, we suspect that the original virus that infected the patient might still be missing in the database because the current closest sequences were circulating in population one and a half years ago, the time was too long, even for a chronic infection. Another hypothesis involves a spillover from humans to animals and spills back from animals to humans; such a process has been proposed to be possible in mink [26]. Interestingly, a recent study proposed that the progenitor of the Omicron variant seemed to have evolved in mice for some time before jumping back into humans [27]. The binding affinity test between the Omicron RBD and animal ACE2 may help to test this hypothesis. A third hypothesis is that the virus split with other variants a long time ago and transmitted cryptically in the population. Since viral genome surveillance is poor in many countries, it is difficult to reject this hypothesis, which again underscores the importance of strengthening viral surveillance on a global scale. Moreover, a hypothesis of acquisition by recombination between different variants is unlikely since the components that make up the Omicron genome could not be found in the current SARS-CoV-2 database, and of course, we cannot reject the possibility that the Omicron genome consists of a combination of components that have not been sequenced. More discussion of the possible origin of the Omicron variant can be found in other studies [28].

Benefiting from the establishment of the viral genome surveillance network and extensive research on the function of viral mutations, it took less than a week to designate the new VOC Omicron since the first identification of its genome, which is much faster than the designation of previous VOCs. However, it will still take several months to verify the risk of the new VOC. There have been over 200,000 new infections per day in the past year. Undoubtedly, we will face more mutant variants in the future, which may result in significant changes in transmissibility, infectivity, and pathogenicity. Unfortunately, it is still impossible to predict the evolutionary direction of the viral genome; hence, we have no hint at what the next VOC will be. To enhance the ability to rapidly respond to the emergence of new VOCs, we should further strengthen genome surveillance on a worldwide scale and develop experimental and computational methods for rapid and high-throughput resolution of mutational functions.

## Materials and methods

### Data collection

The sequencing data were retrieved from SRA database in NCBI (BioProject: PRJNA784038), which was generated by the Network for Genomic Surveillance in South Africa (NGS-SA) [2,6]. In total 211 samples were downloaded on November 30, 2021 (Table S1). The virus lineage was assigned by Pango [29], four samples that cannot be assigned to the Omicron lineage were discarded. All the remaining 207 samples were assigned to Omicron BA.1.

### Quality control and mutation detection

Quality control and adaptor trimming were performed by FASTP [30]. The resultant reads were mapped to Wuhan-Hu-1 (NC_045512.2) using minimap2 (-ax sr) [31]. Primer alignment and trimming were performed by the align_trim function from Artic (https://artic-tools.readthedocs.io/en/latest/commands/#align_trim). The mpileup file and the read count file were generated by SAMtools [32] and Varscan2 [33]. The consensus sequence was obtained using the following criteria: 1) depth ≥ 5-fold; 2) frequency of the major allele ≥ 70%.

### Sequence depth analysis

The sequencing depth was calculated for each nonoverlapping window with a size of 100 bp, except for the last window, which ranged from 29801 to 29880 bp. The fold change of each primer region was calculated by the sum of the depth of all samples in this region divided by the expected value (assuming no differences among regions).

### Identification of epitope regions on the Spike Protein

We downloaded the structures of 182 protein complexes of antibodies that bound to the SARS-CoV-2 *Spike* or its receptor-binding domain (RBD) or N-terminal domain (NTD) from the Protein Data Bank (all structures available before August 8, 2021, www.rcsb.org). The residues in the Spike protein involved in binding to antibodies were identified by a distance of less than 4.5 Å between two counterparts in which van der Waals interactions occur. Deep mutational scanning results were obtained from https://jbloomlab.github.io/SARS2_RBD_Ab_escape_maps/, which includes information on sites in the SARS-CoV-2 RBD where mutations reduce binding by antibodies/sera [22]. The escape score at each position was calculated as the mean of the scores of all antibodies belonging to the same class.

### Construction of the phylogenetic tree

The amino acid sequences were converted from nucleotide sequences using MEGA-X (10.1.8) [34]. Phylogenetic construction was performed by IQ-TREE (1.6.12) [35]. The GTR+F model was used for nucleotide sequences, while the Blosum62 model was used for amino acid sequences.

### TMRCA estimation

The estimation of the time to the most recent common ancestor (TMRCA) and mutation rate was performed by BEAST (2.6.4) [36] using 108 sequences collected between November 13, 2021, and November 23, 2021. The HKY85 nucleotide substitution model and strict molecular clock were used.

### Search for the closest sequences in the database

The distance of two SARS-CoV-2 sequences was represented by the mutation difference, which was calculated by an online tool at National Genomics Data Center, China National Center for Bioinformation https://ngdc.cncb.ac.cn/ncov/online/tool/genome-tracing/?lang=en. Publicly available SARS-CoV-2 sequences were downloaded from the GISAID, NCBI, and RCoV19 databases (November 1, 2021) [1,37].

### Calculation of linkage disequilibrium

The *r*^2^ statistic was used to measure the strength of the linkage disequilibrium between each pair of mutations [38]. The calculation of linkage disequilibrium was based on all unique haplotypes from the public database.

## CRediT author statement

**Wentai Ma:** Methodology, Formal analysis, and Writing. **Jing Yang:** Methodology, Formal analysis, and Writing. **Haoyi Fu:** Methodology, Formal analysis. **Chao Su:** Methodology, Formal analysis. **Caixia Yu:** Resources. **Qihui Wang:** Methodology. **Ana Tereza Ribeiro de Vasconcelos:** Resources, Writing. **Georgii A. Bazykin:** Resources, Writing. **Yiming Bao:** Methodology, Writing. **Mingkun Li:** Conceptualization, Methodology, Supervision, Writing. All authors have read and approved the final manuscript.

## Acknowledgments

We thank Dr. Jennifer Giandhari, Dr. Eduan Wilkinson, and Dr. Tulio de Oliveira from Centre for Epidemic Response and Innovation (CERI), Stellenbosch University for sharing the data and workflow, GISAID and associated laboratories and researchers for the shared sequence information, Dr. Aiping Wu from Suzhou Institute of Systems Medicine, Chinese Academy of Medical Sciences, for help in the phylogenetic tree analysis. This study was funded by the National Natural Science Foundation of China (Grant No. 82161148009), the Strategic Priority Research Program of Chinese Academy of Sciences, China (Grant No. XDB38030400), The Capital Health Development and Research of Special (Grant No. 2021-1G-3012), Conselho Nacional de desenvolvimento Cientifico e Tecnológico (CNPq) - NGS-BRICS - n°: 440931/2020-7, and Russian Foundation for Basic Research (RFBR) (Grant No. 20-54-80014.

## Supplementary material

**Figure S1. The correlation of the major allele frequency between the Illumina protocol and the GridION protocol**.

Each dot represents a mutation (major allele frequency >= 70%).

**Figure S2. The distribution of the Omicron mutations on the structure of the Spike protein (left) and the RBD region (right)**.

**Table S1. The information of the data used in the study**.

