## Supplementary figures and images for "Genomic Perspectives on the Emerging SARS-CoV-2 Omicron Variant"

### Figure S1.tif

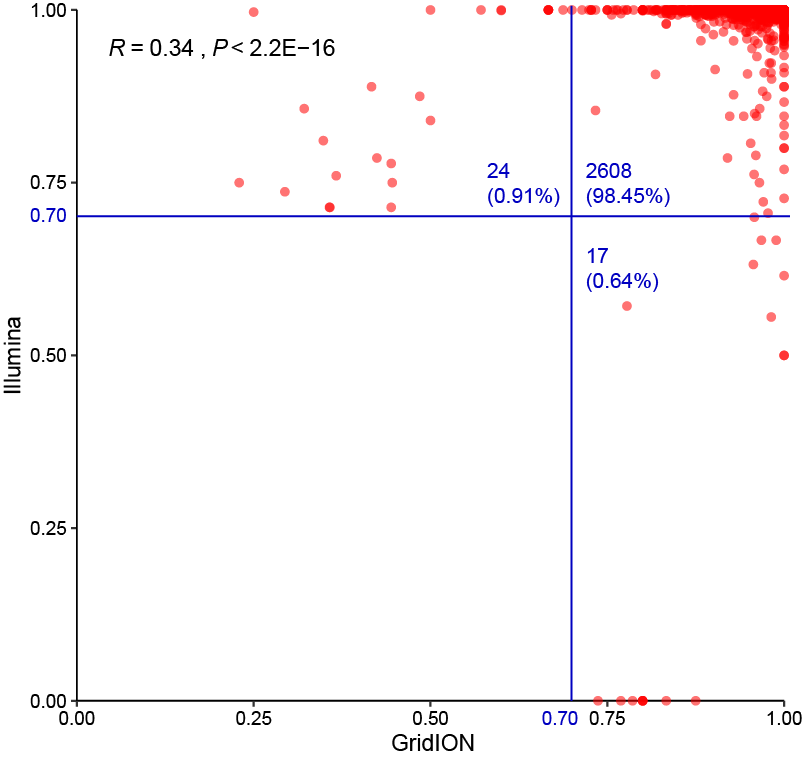

### Figure S2.tif

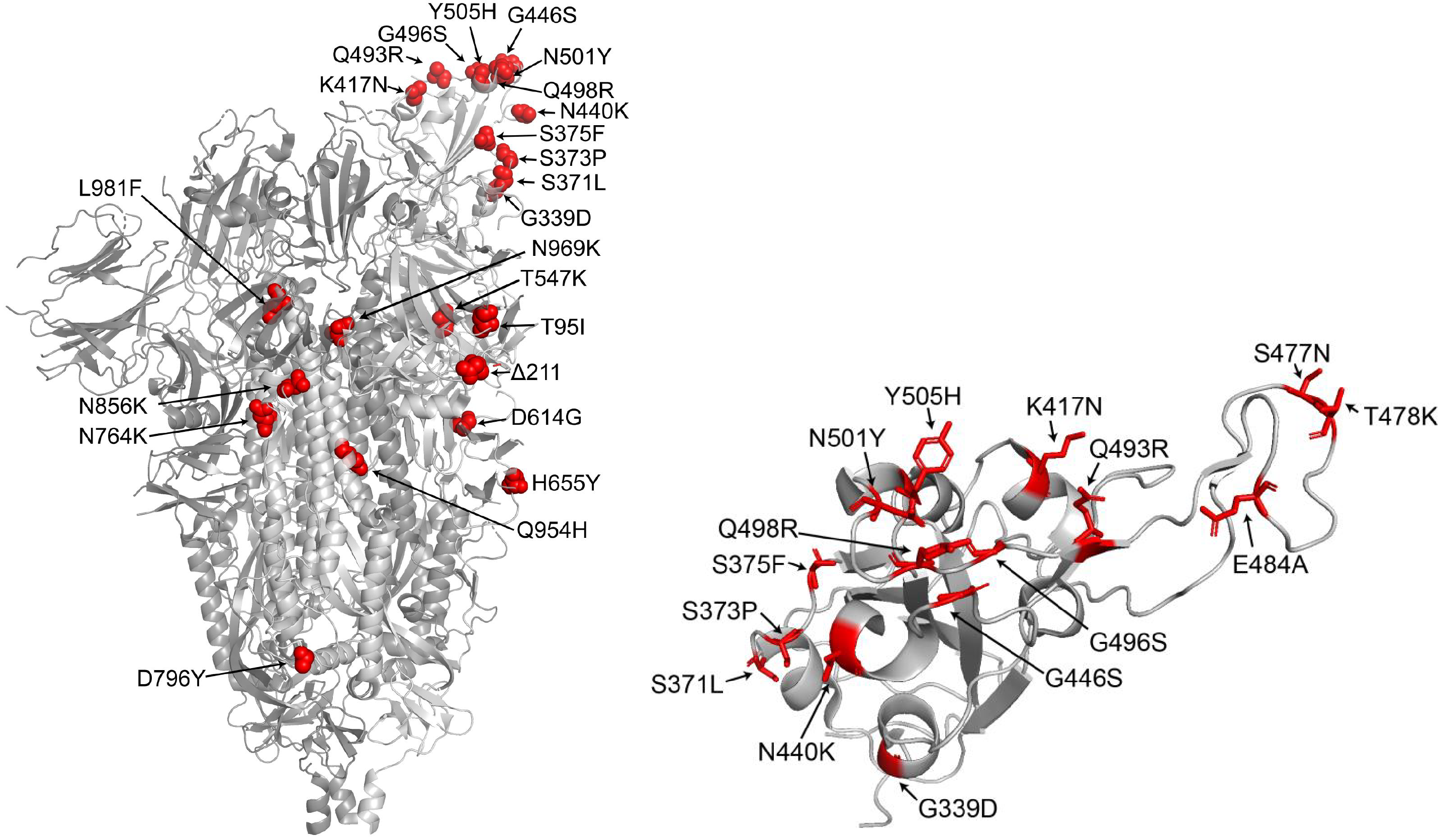
